# Intrinsic Resistance of the Hippocampal CA2 Subfield to Neuroinflammation After Status Epilepticus

**DOI:** 10.1101/2025.07.29.667443

**Authors:** Bruno Ponciano Da silva, Edna Cristina Santos Franco, Silene Maria Araújo de Lima

## Abstract

**Objective:** To determine the spatiotemporal patterns of pro-inflammatory (IL-1beta) and anti-inflammatory (IL-10) cytokines in hippocampal subfields, focusing on the CA2 region, using a pilocarpine-induced status epilepticus (SE) model.

**Methods:** Status epilepticus was induced in adult male Wistar rats with pilocarpine. Animals were divided into control, 1-day post-SE and 7-day post-SE groups (n = 5, 3 and 3). Hippocampi were processed for immunohistochemistry using antibodies against IL-1beta, IL-10, NeuN (mature neurons), and PCP4 (CA2 marker). Microglial activation states (M1/M2) were inferred from cytokine profiles: sustained IL-1beta expression indicated a pro-inflammatory milieu (M1), whereas declining IL-1beta in the presence of IL-10 suggested an anti-inflammatory or reparative state (M2).

**Results:** The CA2 region exhibited IL-1beta immunoreactivity at 1 day post-SE, which decreased by day 7, while CA3 maintained elevated IL-1beta levels. Anti-IL-10 immunostaining was prominent across hippocampal subregions in the control group and 1-day SE group but was absent by day 7 in all regions. NeuN staining revealed limited neuronal death in CA2 at 1 day post-SE, with substantial loss across CA1, CA3, and CA4 by day 7.

**Significance:** The CA2 subfield appears relatively protected from sustained inflammation and neuronal loss, likely owing to unique microglial responses and structural features such as perineuronal nets. These findings highlight microglial polarization as a potential determinant of subfield vulnerability in temporal lobe epilepsy and support further investigation of glial-targeted therapies.

## Introduction

Temporal lobe epilepsy (TLE) is the most common form of focal epilepsy in adults and is frequently drug-resistant (Fisher et al., 2005; Scheffer et al., 2017). One of its main neuropathological substrates is hippocampal sclerosis (HS), characterized by selective neuronal loss, gliosis, and circuit reorganization within specific hippocampal subfields, particularly CA1, CA3, CA4, and the dentate gyrus (Blümcke et al., 2013; Thom, 2014). Despite being surrounded by vulnerable regions, the CA2 subfield consistently exhibits structural preservation and relative resistance to epileptic injury, suggesting a potentially neuroprotective microenvironment and a unique role in the epileptic hippocampal network (Steve et al., 2014; Kilias et al., 2023).

The pathophysiology of TLE is intricately linked to neuroinflammation. Following a precipitating event such as status epilepticus (SE), the central nervous system activates immune-like responses, leading to the release of cytokines and chemokines predominantly by glial cells, including microglia, and neuronal cells (Vezzani et al., 2011; Devinsky et al., 2013). Microglia, as resident immune cells of the brain, undergo rapid activation and can exhibit distinct functional states: pro-inflammatory (M1) and anti-inflammatory or reparative (M2), each characterized by specific cytokine profiles. Among the key mediators of this response are interleukin-1 beta (IL-1β), a pro-inflammatory cytokine predominantly released by activated microglia (M1 phenotype) known to exacerbate excitotoxicity and neuronal hyperexcitability, and interleukin-10 (IL-10), an anti-inflammatory cytokine typically associated with the reparative microglial phenotype (M2), which modulates immune activation and promotes neuroprotection (Vezzani et al., 2002; Maroso et al., 2011). An imbalance between these mediators, potentially driven by microglial polarization, has been associated with the initiation and perpetuation of epileptogenesis. However, the spatial and temporal expression of IL-1β and IL-10, and their indirect implications for microglial activity across hippocampal subfields—particularly within CA2—remain poorly understood.

Previous histopathological studies and experimental models have demonstrated that CA2 possesses unique molecular and structural features, including robust perineuronal nets, low seizure susceptibility, and expression of the specific neuronal marker Purkinje cell protein 4 (PCP4) (San Antonio et al., 2014; Carstens et al., 2016). Such characteristics might influence the local microenvironment and microglial responses, raising the hypothesis that CA2 is equipped with endogenous mechanisms involving distinct microglial cytokine profiles that shield it from excitotoxic and inflammatory damage. Yet, little is known about how inflammatory cascades and microglial dynamics operate within CA2 during the early stages of epileptogenesis.

In the context of mesial temporal lobe epilepsy (MTLE), where HS and inflammation coexist, investigating cytokine dynamics and their indirect association with microglial polarization in CA2 may provide critical insights into the mechanisms underpinning regional vulnerability or resistance. Furthermore, identifying whether CA2 is protected by anti-inflammatory microglial responses—or simply spared due to lack of engagement—could inform targeted interventions that leverage neuroprotective processes in adjacent, more vulnerable areas.

This study aimed to characterize the expression patterns of IL-1β and IL-10 in hippocampal subfields during the early phases following SE induced by pilocarpine in adult rats. Using immunohistochemical labeling with PCP4 to delineate the CA2 subfield, we analyzed neuronal integrity and cytokine distribution at two time points (1 and 7 days post-SE). We hypothesize that CA2 displays a distinct cytokine profile indicative of unique microglial dynamics, marked by transient pro-inflammatory signaling and reduced neurodegeneration compared to neighboring subfields, which may underlie its relative preservation in TLE.

Our findings contribute to a more nuanced understanding of CA2’s role in epileptogenesis, indirectly highlight microglial polarization as a potential determinant of subfield vulnerability, and emphasize the importance of spatially resolved inflammatory analyses in experimental epilepsy models. This work may pave the way for future therapeutic approaches that aim to mimic or enhance the protective environment of CA2 to mitigate hippocampal damage and seizure propagation.

## Materials and Methods

### Experimental Animals

Thirty adult male Wistar rats (250–300 g) were initially considered; however, only 11 animals were ultimately included in the final analysis (see Results). All animals were bred at the Central Animal Facility of the Federal University of Pará (UFPA) and housed individually under controlled environmental conditions (22–25 °C; 12/12 h light/dark cycle), with food and water provided ad libitum and regular ectoparasite control. All experimental procedures were approved by the UFPA Animal Research Ethics Committee (CEUA Nº 3347221217) and complied with international standards for animal care and use.

### Induction of Status Epilepticus

To induce status epilepticus (SE), animals were first pretreated with methyl-scopolamine (1 mg/kg, s.c.) 30 minutes before pilocarpine to minimize peripheral cholinergic effects. Pilocarpine hydrochloride was administered intraperitoneally at incremental doses of 100, 200, and 360 mg/kg diluted in sterile saline. Seizure severity was assessed behaviorally using the Racine scale (1972). Only animals reaching stages 3–5 within one hour post-injection were included as having developed SE and were treated with diazepam (10 mg/kg, i.p.) to interrupt seizures and reduce mortality.

The Racine scale was used to classify seizure behavior as follows:

- Stage 0: No behavioral change
- Stage 1: Sudden facial and snout movements
- Stage 2: Head nodding
- Stage 3: Forelimb clonus
- Stage 4: Rearing
- Stage 5: Rearing and falling with generalized tonic-clonic seizures

### Experimental Groups

The rats were assigned to three experimental groups:

- **Control group (n = 5):** received saline and were euthanized 7 days later.
- **SE-1D group (n = 3):** subjected to pilocarpine-induced SE and euthanized 1 day post-SE.
- **SE-7D group (n = 3):** subjected to pilocarpine-induced SE and euthanized 7 days post-SE.

Animals were monitored and housed during the survival period at the Laboratory of Experimental Neuroprotection and Neuroregeneration, a facility accredited by the Brazilian National Council for Animal Experimentation (CONCEA).

### Perfusion and Histological Preparation

At the defined survival endpoints (1 or 7 days), animals were anesthetized with ketamine (72 mg/kg) and xylazine (9 mg/kg), both administered intraperitoneally. Adequate anesthesia was confirmed by the absence of corneal and withdrawal reflexes. Rats were transcardially perfused via the left ventricle with 500 mL of heparinized saline (0.9%), followed by 500 mL of 4% paraformaldehyde (PFA) in 0.1 M phosphate buffer (PB, pH 7.2–7.4).

Brains were post-fixed for 24 hours and cryoprotected in increasing concentrations of sucrose diluted in a solution of glycerol and 0.05 M PB. Tissues were embedded in OCT (Tissue-Tek, Sakura), frozen, and coronally sectioned (30 µm) using a Leica CM3050 cryostat. Sections were stored in cryoprotectant at −20 °C.

### Nissl Staining

For histopathological analysis, selected sections were stained with 0.1% thionin (Nissl staining) to detect Nissl bodies in the neuronal cytoplasm. The presence or absence of these structures served as an indicator of neuronal viability or chromatolysis.

### Immunohistochemistry

Sections were washed in distilled water and PBS, followed by antigen retrieval in 90 °C citrate buffer for 20 minutes. Permeabilization was performed using saponin solution for 10 minutes. After blocking with 10% bovine serum albumin (BSA) for 30 minutes, slices were incubated overnight at 4 °C with the primary antibodies listed below.

The next day, sections were washed and incubated with biotinylated secondary antibodies (anti-mouse and anti-rabbit, Dako REF K0675) for 30 minutes at 37 °C, followed by streptavidin-HRP for 30 minutes. Visualization was achieved using 3,3’-diaminobenzidine (DAB) as the chromogen. In some cases, hematoxylin counterstaining was performed. Sections were dehydrated in a graded ethanol series, cleared in xylene, and coverslipped with Permount (Fisher Scientific).

### Primary antibodies used

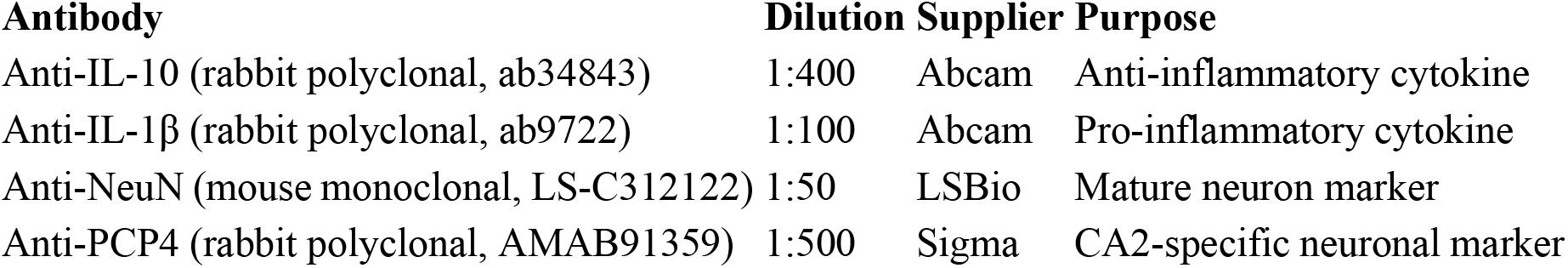

Although microglia were not directly visualized using specific microglial markers in this study, the analysis of pro-inflammatory (IL-1β) and anti-inflammatory (IL-10) cytokines served as indirect indicators of microglial functional states (M1/M2 phenotypes), as previously established in the literature (Vezzani et al., 2011; Orihuela et al., 2016).

### Image Acquisition and Qualitative Analysis

Histological analysis and photodocumentation were performed using a Carl Zeiss Imager.Z1 microscope equipped with AxioCam Hrc and AxioVision software. The CA2 subfield was identified via PCP4 immunolabeling. Photomicrographs were qualitatively analyzed based on hippocampal cytoarchitecture and compared to known features described in the literature for mesial temporal lobe epilepsy.

**Note:** Due to logistical constraints during the COVID-19 pandemic and high mortality rates following SE induction, this study was conducted with a reduced number of animals. At the time, resource limitations precluded expanding the sample size or performing quantitative analyses. As such, our histological and immunohistochemical evaluations were limited to a qualitative approach. A follow-up study with an increased number of animals and statistical power is planned to validate and extend these findings through quantitative and stereological methods.

## Results

### Behavioral Observations and Mortality in the Pilocarpine Model

Of the 30 adult male Wistar rats initially available, five were assigned to the control group and received saline, while 25 were subjected to the pilocarpine-induced status epilepticus (SE) protocol. Out of the 25 experimental animals, only six survived the entire SE induction protocol (methyl-scopolamine, pilocarpine, and diazepam) and the subsequent survival periods of 1 or 7 days. Nineteen animals died at various stages of the protocol, which lasted approximately two hours in total. Behavioral seizure severity was assessed using the Racine scale (1972), with animals reaching stages 3–5 considered to have entered SE and included in the study.

### Hippocampal Integrity and Neuronal Loss – NeuN Staining

Anti-NeuN immunostaining was used to evaluate neuronal integrity. In control animals, NeuN expression was robust and widespread throughout the hippocampal subfields, including CA1, CA3, the dentate gyrus (DG), and hilus, with preserved cytoarchitecture (Figure 5A, B).

**Figure 5.**
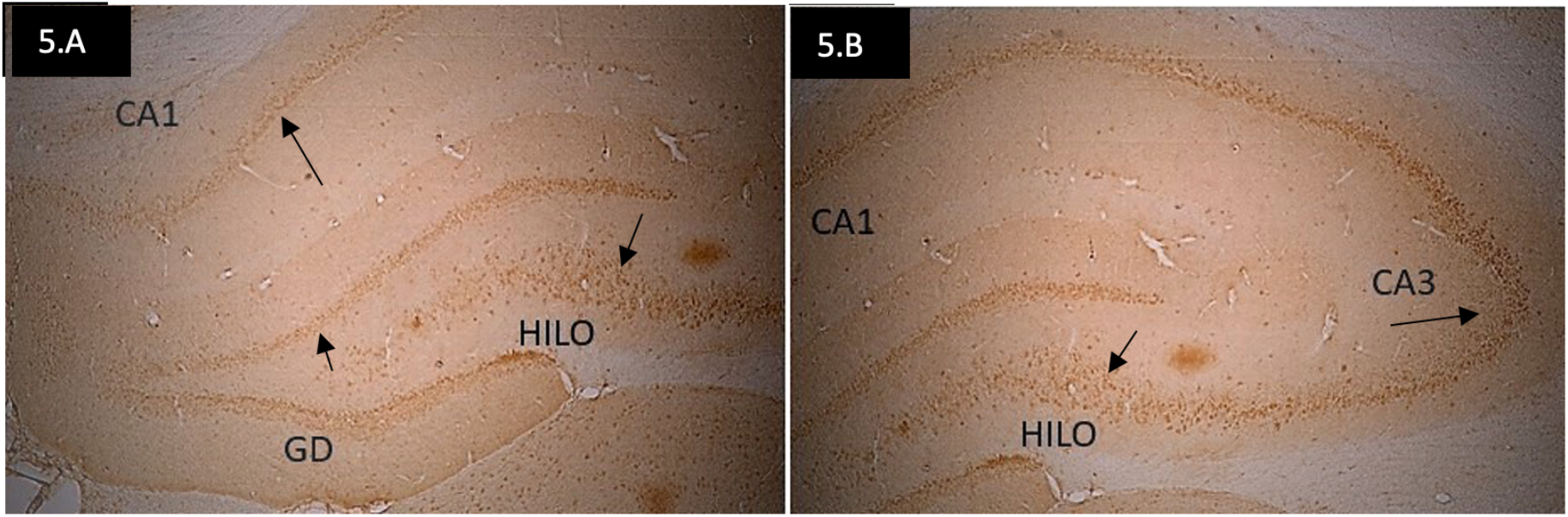
Photomicrographs of hippocampi from control animals stained with anti-NeuN. Panels A and B display the left and right hemispheres of the same hippocampus, showing preserved neuronal layers in CA1, CA3, DG, and hilus. Arrows indicate NeuN-positive mature neurons (objective: 5×).

In SE animals, NeuN immunoreactivity was reduced, particularly in the CA3 region after 1 day (Figure 6C), suggesting neuronal loss. At 7 days post-SE, there was a pronounced reduction in NeuN labeling across CA1, CA2, CA3, and the hilus, indicating progressive neuronal degeneration (Figure 7C, D), in contrast to preserved labeling in controls (Figure 7A, B).

**Figure 6.**
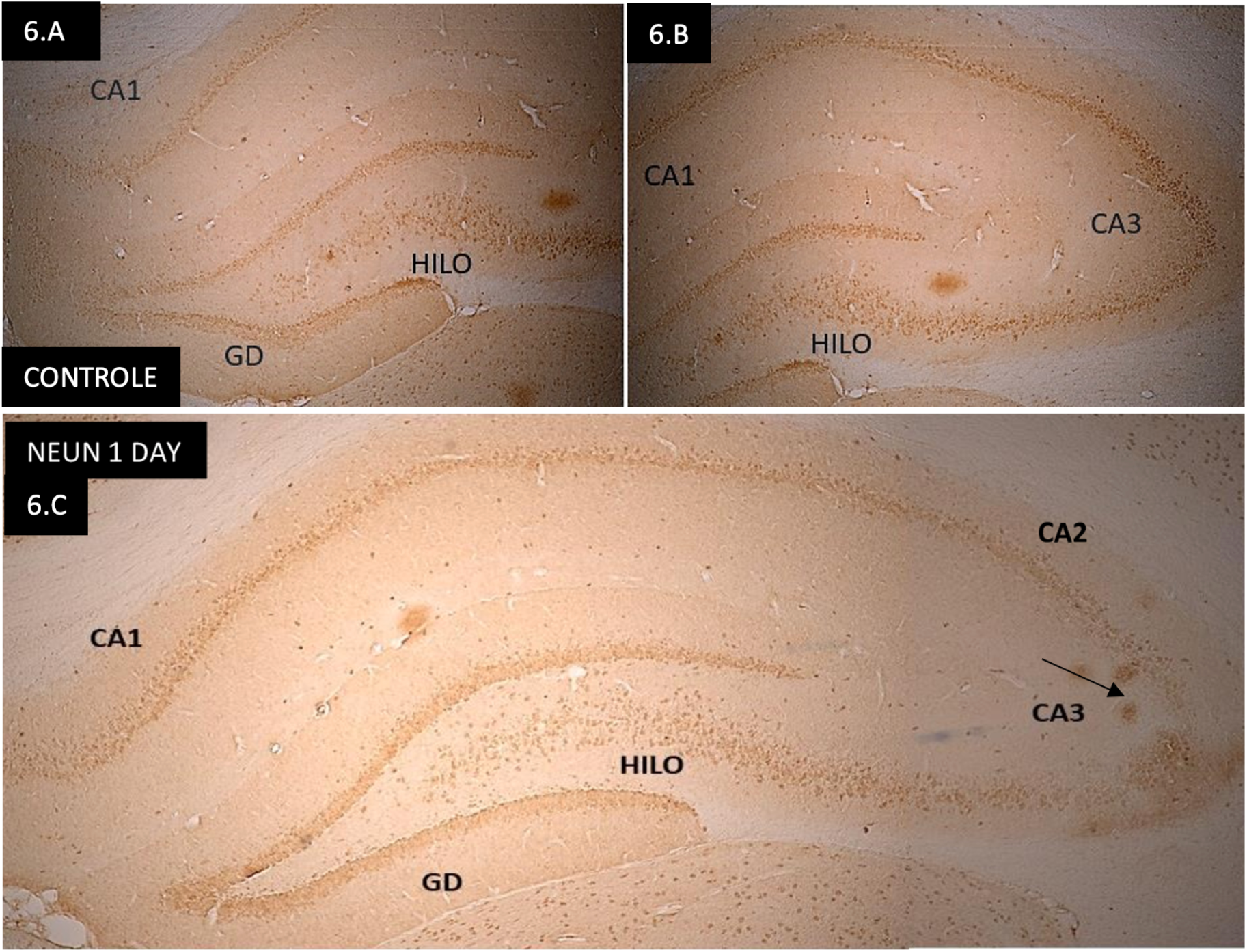
NeuN-stained hippocampal sections. Panels A and B: control hippocampus showing preserved NeuN labeling in CA1, CA2, CA3, DG, and hilus. Panel C: SE animal (1 day post-SE) showing reduced NeuN immunoreactivity in CA3 (objective: 5×).

**Figure 7.**
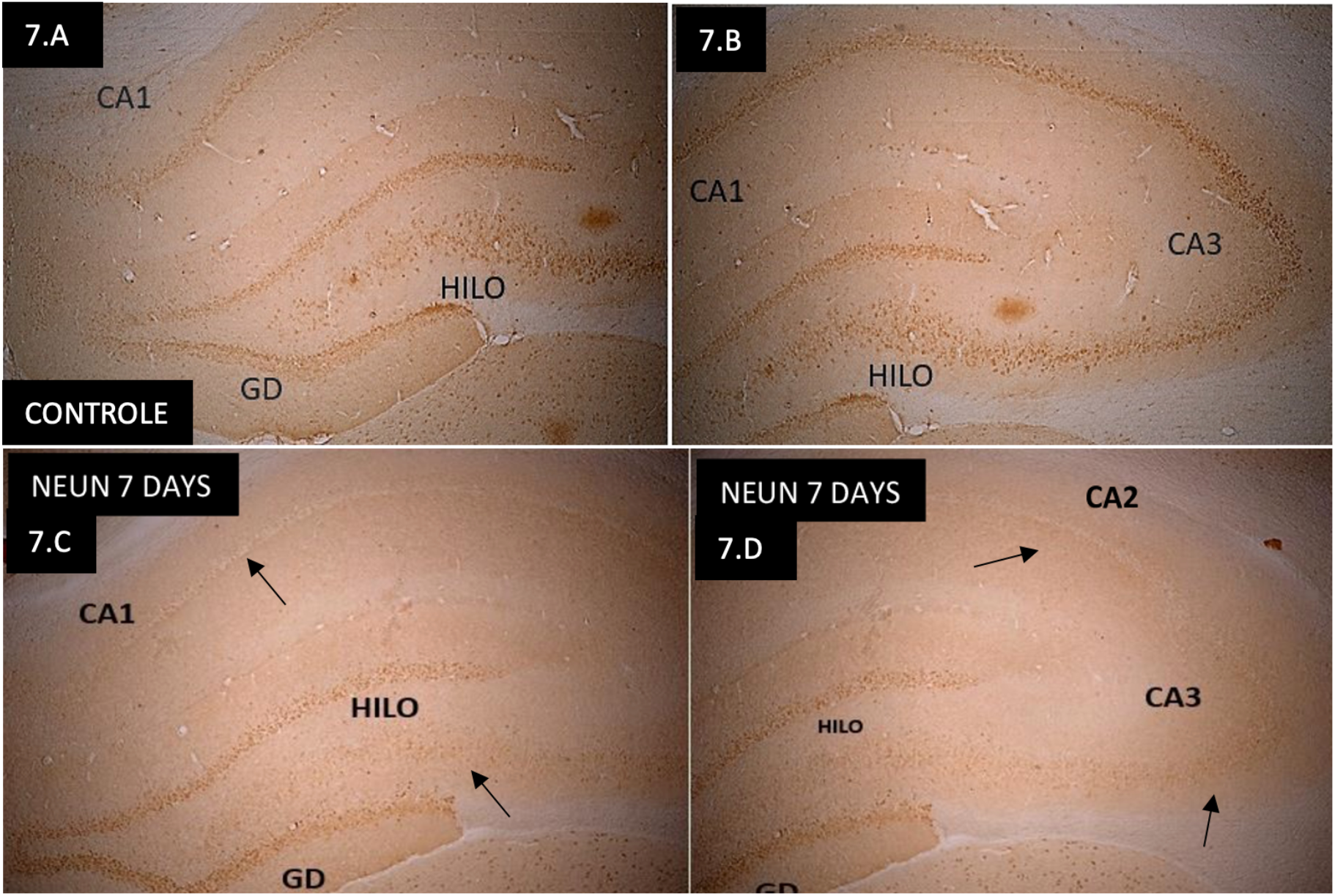
NeuN immunostaining in hippocampi. Panels A and B: control animals with no neuronal loss. Panels C and D: SE animals at 7 days post-SE showing marked reduction of NeuN labeling in CA1, CA2, CA3, and hilus. Arrows indicate regions of neuronal loss (objective: 5×).

### CA2 Identification via PCP4 Labeling

The CA2 region was clearly delineated using PCP4 immunolabeling. In control animals, anti-PCP4 distinctly labeled CA2 pyramidal neurons, allowing precise differentiation from adjacent subfields CA1 and CA3 (Figure 8A–C).

**Figure 8.**
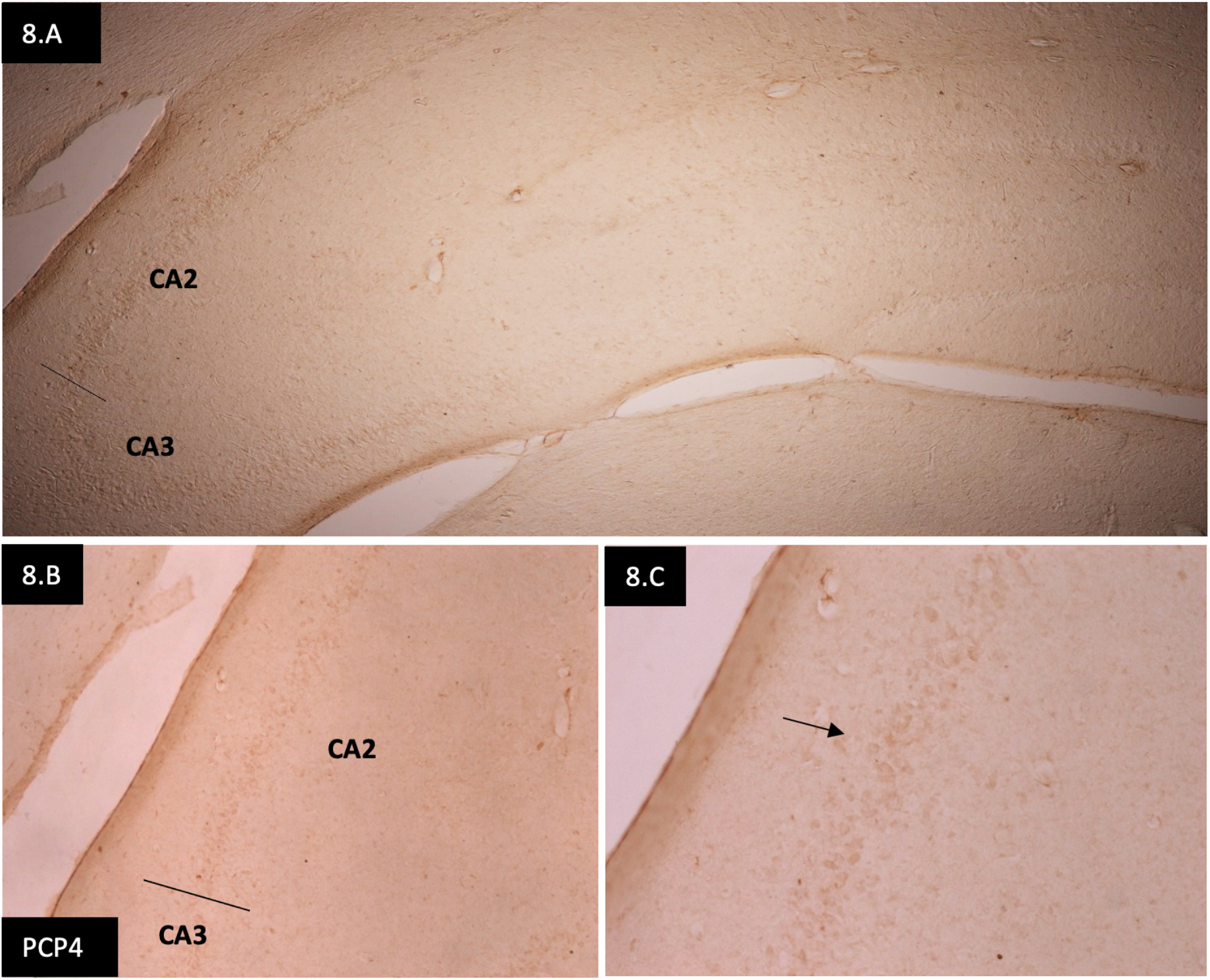
PCP4 immunolabeling in control hippocampi. Panel A: low magnification overview of the hippocampus. Panel B: demarcation of the CA2/CA3 boundary (arrow). Panel C: high magnification showing pyramidal neurons of CA2 (objectives: A = 5×; B = 10×; C = 20×).

**Figure 9.**
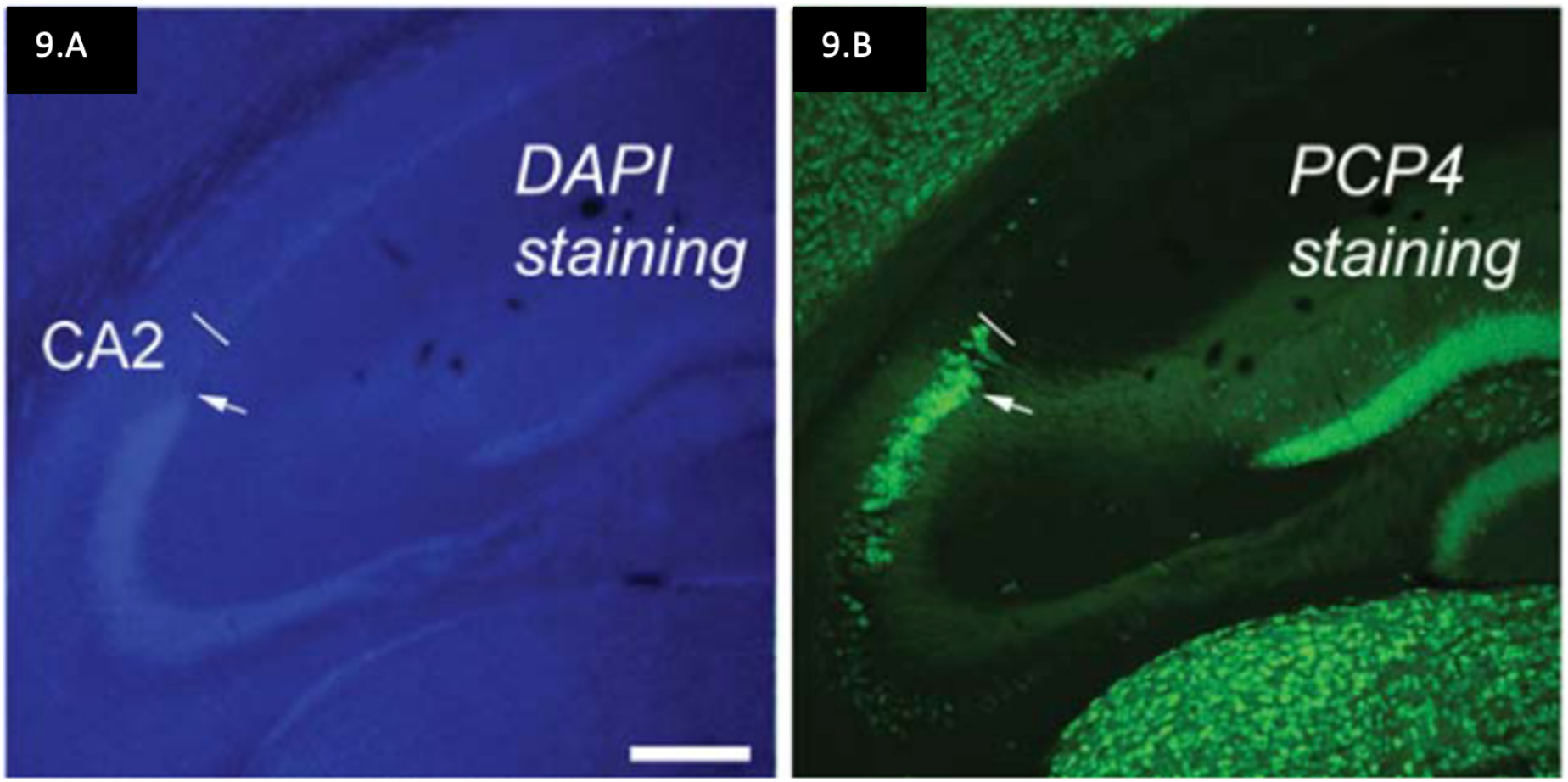
Reference data from literature showing PCP4 immunoreactivity in adult mouse CA2. Panel A: DAPI nuclear stain. Panel B: PCP4 stain with boundaries of CA2 clearly indicated (adapted from San Antonio et al., 2014).

### Anti-Inflammatory Cytokine Expression – IL-10

IL-10 expression was assessed to explore the anti-inflammatory profile of the hippocampus. Control animals showed widespread IL-10 immunoreactivity throughout all hippocampal subregions (Figure 10A, B), consistent with a role in maintaining homeostasis. However, in SE animals, IL-10 immunoreactivity decreased markedly over time. At 1 day post-SE, faint IL-10 labeling was seen in CA3 and DG (Figure 10C, D), while by day 7, IL-10 labeling was absent in all regions examined (Figure 10E, F).

**Figure 10.**
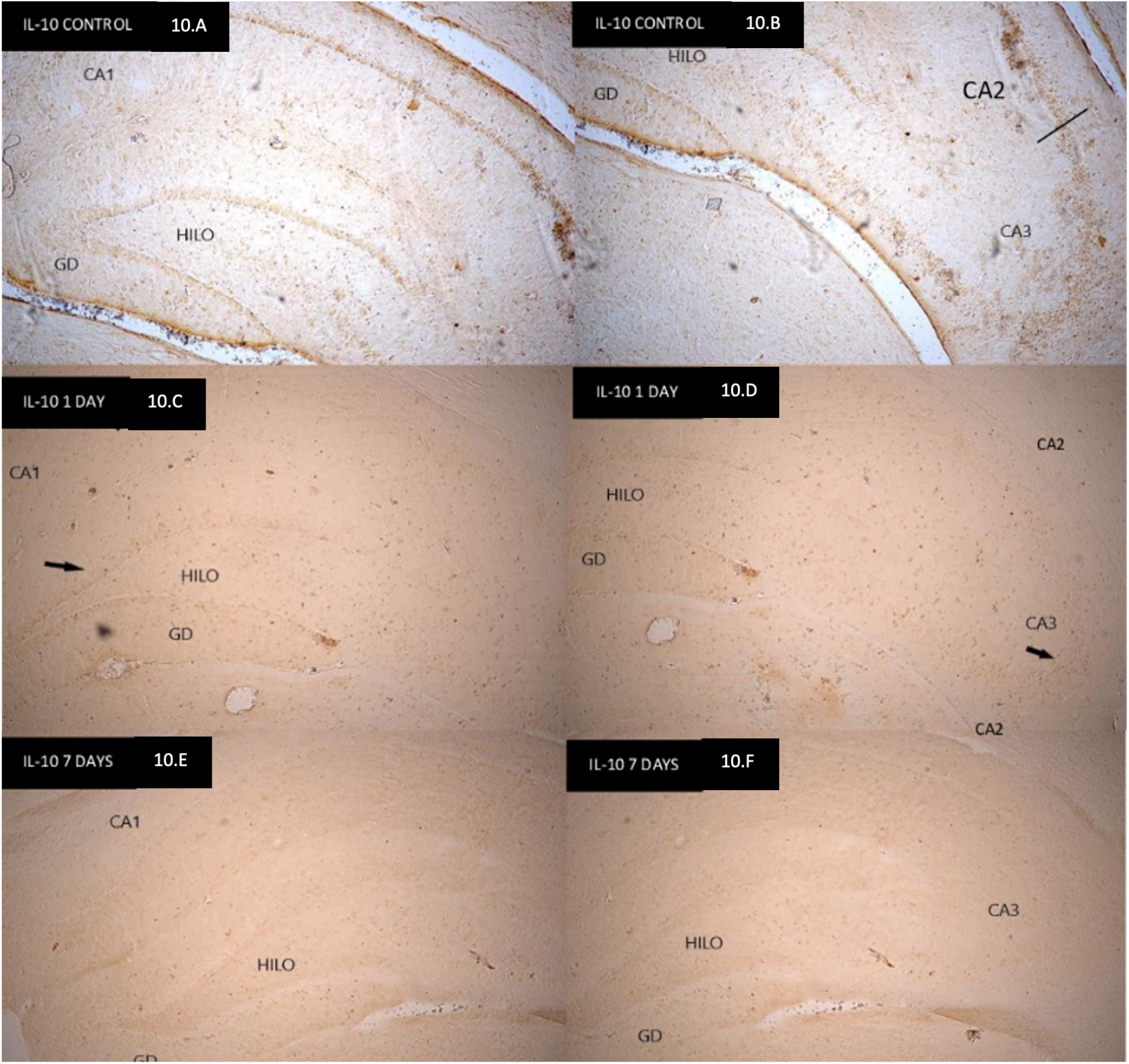
IL-10 immunolabeling in hippocampal sections. Panels A and B: control animals showing widespread labeling in all subregions. Panels C and D: SE animals (1 day post-SE) with reduced IL-10 in CA3 and DG. Panels E and F: SE animals (7 days post-SE) with no detectable IL-10 labeling (objective: 5×).

The progressive decline in IL-10 expression after SE further supports a potential change in microglial functional state, indicating a decreased anti-inflammatory response over time.

### Pro-Inflammatory Cytokine Expression – IL-1β

IL-1β immunostaining was used to assess pro-inflammatory response. At 1 day post-SE, IL-1β was prominently expressed in CA2 and CA3 (Figure 11A), indicating activation of neuroinflammatory pathways in these regions. At 7 days post-SE, IL-1β expression had diminished in CA2 but remained evident in CA3 (Figure 11B), suggesting a spatial and temporal shift in inflammatory activity.

**Figure 11.**
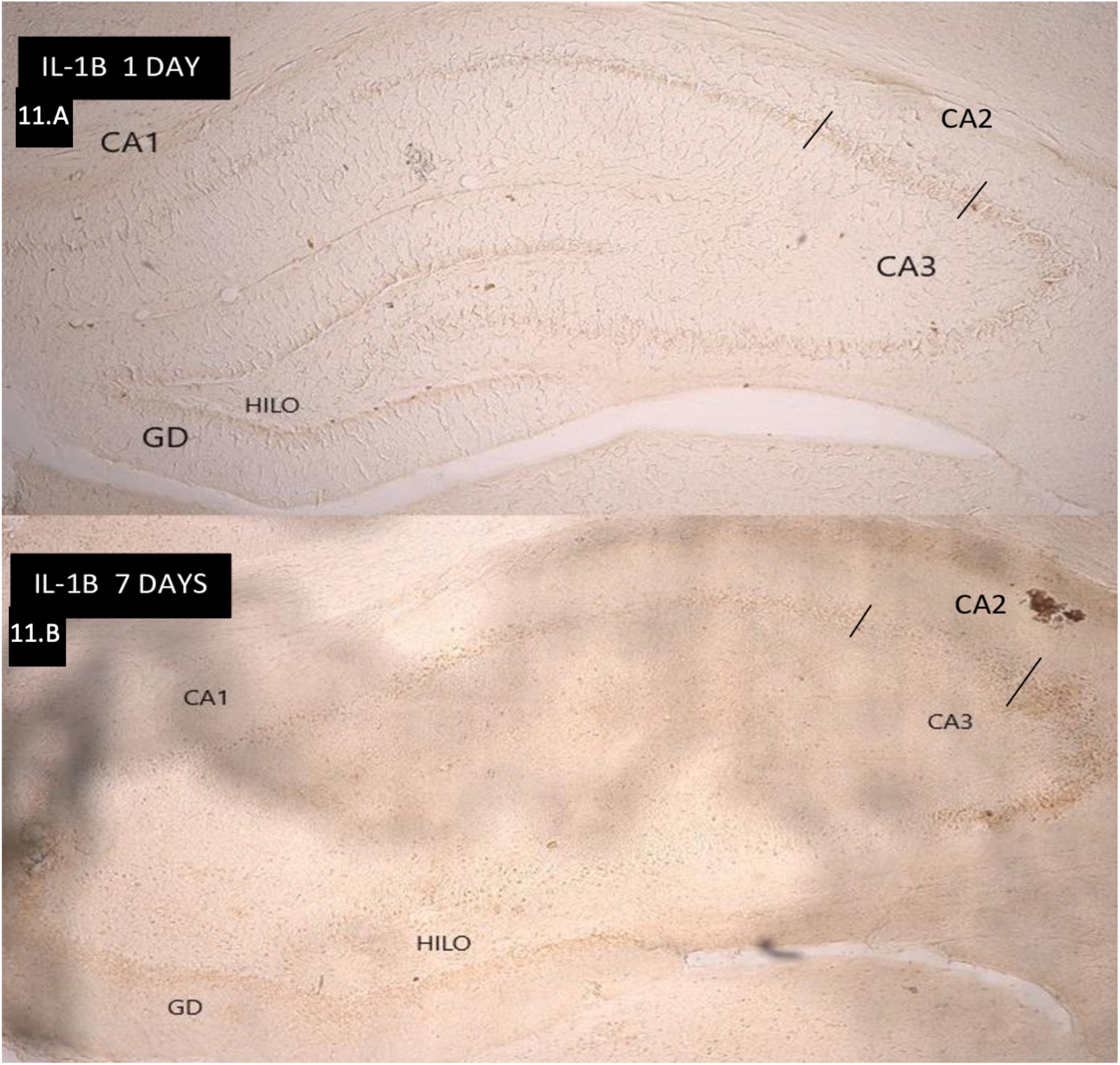
IL-1β immunostaining in hippocampal tissue. Panel A: SE animal (1 day post-SE) showing strong IL-1β expression in CA2 and CA3. Panel B: SE animal (7 days post-SE) showing reduced labeling in CA2 but persistent labeling in CA3 (objective: 5×).

A reduction in IL-1β immunoreactivity in the CA2 region at 7 days post-SE may suggest a shift in microglial polarization from a pro-inflammatory M1 state towards an anti-inflammatory or reparative M2 state. Although microglia were not directly labeled, these cytokine expression patterns indirectly indicate differential microglial responses across hippocampal subfields.

### Qualitative Nature and Sample Limitations

This study was based on qualitative histological analysis due to the limited number of animals surviving the SE protocol and the constraints imposed by the COVID-19 pandemic. Although this limits statistical generalization, the data provide valuable insight into cytokine dynamics and neuronal vulnerability across hippocampal subfields, particularly CA2. Future studies with increased sample sizes and quantitative approaches are planned to validate and extend these findings.

## Discussion

This study investigated the temporal dynamics of neuronal loss and cytokine expression in hippocampal subfields following pilocarpine-induced status epilepticus (SE), with a particular focus on the CA2 region. Qualitative immunohistochemical analyses revealed differential vulnerability across subfields and point to CA2 as a potentially unique site of neuroinflammatory regulation in the context of epileptogenesis.

NeuN staining demonstrated progressive neuronal loss, with greater reduction in NeuN-positive cells in CA1, CA2, CA3, and the hilus at 7 days post-SE compared to 1 day. These results are consistent with the literature describing selective vulnerability of hippocampal neurons after SE (Thom, 2014; Steve et al., 2014). The observed reduction of NeuN immunoreactivity in CA2 contrasts with earlier assumptions that this subfield is largely spared in temporal lobe epilepsy (TLE); however, recent studies have reported variable patterns of CA2 involvement depending on seizure severity and duration (Kilias et al., 2023).

The inflammatory profile observed in this study supports region- and time-dependent modulation of cytokine responses. In control animals, IL-10 was diffusely expressed throughout the hippocampus, confirming its role in maintaining basal anti-inflammatory tone. This is in agreement with findings that IL-10 is constitutively expressed in healthy brain tissue and modulates neuroimmune interactions. However, IL-10 expression decreased sharply by 1 day post-SE and was virtually absent by 7 days, suggesting an insufficient or exhausted anti-inflammatory response following prolonged seizures. This may be a key factor contributing to the chronicity of inflammation and progression of neuronal damage in epileptic tissue. This pattern aligns with previous findings showing that deficient IL-10 expression in the hippocampus correlates with increased neurodegeneration, as observed in a neurotoxic model of TMT-induced injury (Kamaltdinova et al., 2021).

Given the decline in IL-10 immunoreactivity and the apparent resilience of CA2, future studies should consider the involvement of other anti-inflammatory cytokines such as IL-4, IL-11, or IL-13, which may contribute to localized neuroprotection through distinct signaling cascades.

Conversely, IL-1β was strongly expressed in CA2 and CA3 at 1 day post-SE, but its expression declined in CA2 while persisting in CA3 at 7 days. This spatiotemporal pattern indicates a potential shift of inflammatory burden from CA2 to CA3, and suggests that CA2 might engage early anti-inflammatory or structural defense mechanisms limiting prolonged cytokine activation. Prior studies have demonstrated that IL-1β plays a pivotal role in mediating seizure-induced neurotoxicity and epileptogenesis, particularly through its downstream effects on COX-2 and prostaglandin signaling pathways (Friedman & Dingledine, 2011; Rojas et al., 2014; Vezzani et al., 2011). In particular, activation of the EP2 receptor by PGE2 has been shown to promote hippocampal inflammation and neuronal death, while systemic inhibition of EP2 reduced mortality, brain inflammation, and neurodegeneration in pilocarpine models of epilepsy (Jiang et al., 2012; Jiang et al., 2013).

Additionally, the observed temporal dynamics of IL-1β and IL-10 indirectly reflect microglial polarization states, supporting the hypothesis of an initial pro-inflammatory (M1-like) activation followed by a potential shift toward an anti-inflammatory or reparative (M2-like) phenotype. Although IL-10 expression declined progressively and was absent at 7 days post-SE, this does not necessarily indicate a cessation of M2-associated activity. Instead, it is plausible that other anti-inflammatory cytokines—such as IL-4, IL-13, or TGF-β—may become predominant at later stages, maintaining the anti-inflammatory tone through alternative signaling. Although microglia were not directly labeled, these cytokine patterns support the relevance of microglial polarization in shaping subfield-specific vulnerability and resilience in epileptogenesis.

The anatomical localization of CA2 using PCP4 immunolabeling was critical for accurate subfield analysis. PCP4 is a well-established marker for CA2 pyramidal neurons and allows precise distinction from CA1 and CA3, which is essential in studies investigating selective vulnerability and inflammation. Our findings confirm that the CA2 region can be reliably identified using PCP4 in both control and post-SE tissue, consistent with previous reports (San Antonio et al., 2014).

The potential for CA2 to exhibit relative resistance to sustained inflammatory damage may be related to unique structural features such as perineuronal nets (PNNs), which are enriched in this subregion and known to modulate synaptic plasticity and excitability (Carstens et al., 2016; Alpár et al., 2006; Bitanihirwe & Woo, 2015). These networks may dampen excessive excitability and act as barriers to the spread of inflammatory signaling, providing structural support that could explain the observed transient IL-1β immunoreactivity in CA2.

A major limitation of the present study is the reliance on qualitative data due to the small sample size and high mortality rate associated with the pilocarpine model. These constraints were further compounded by logistical challenges during the COVID-19 pandemic. Despite this, the consistency of observed patterns across samples lends preliminary support to our hypotheses. Future work will include quantitative immunohistochemical and molecular analyses to validate these findings and investigate additional cytokines and structural markers implicated in hippocampal resilience.

Overall, this study contributes to a growing body of literature suggesting that the CA2 region may play a unique role in the inflammatory landscape of the epileptic hippocampus. Further research into the molecular underpinnings of CA2’s differential response to SE could reveal novel targets for anti-inflammatory or neuroprotective interventions in temporal lobe epilepsy.

## Conclusion

This study reveals temporal and spatial differences in cytokine expression and neuronal survival in the hippocampus post-SE. Cytokine patterns indirectly suggest that subfield-specific inflammatory responses may involve shifts in microglial polarization, even as IL-10 expression declines over time. The CA2 region exhibits a unique inflammatory profile and resistance to neuronal loss, offering insights into potential neuroprotective mechanisms relevant to TLE.

## Acknowledgments

This work was supported by CAPES-CNPQ. The authors thank Evantro Chagas Institute and Federal University of Pará for their support in conducting the experiments.

